## Supplementary Figures for "HyperChIP for identifying hypervariable signals across ChIP/ATAC-seq samples"

### Supplementary Figure1

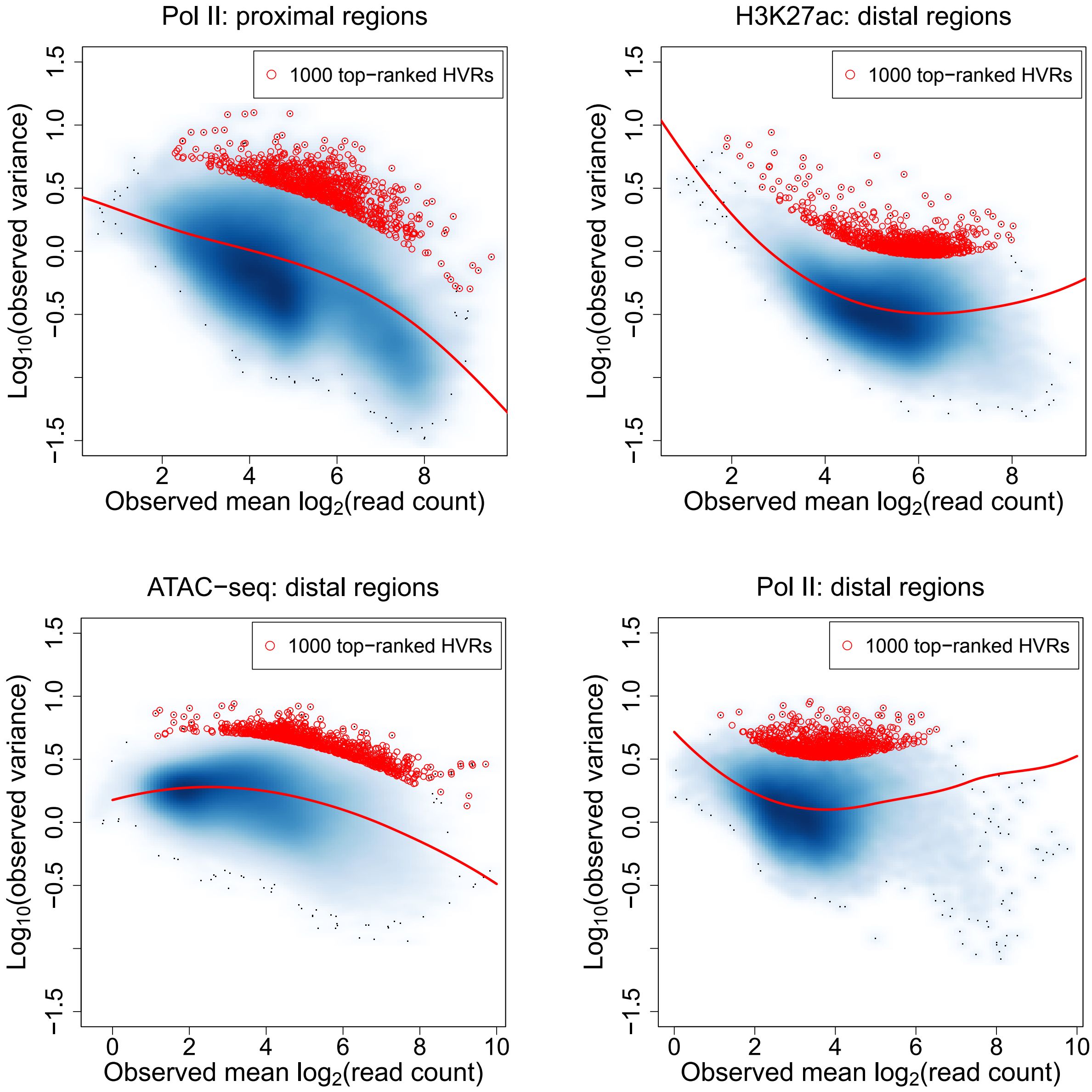

**Supplementary Figure 1. Scatter plots showing various mean-variance trends associated with different data sets.** Variance is shown at the $\log_{10}$ scale. Red lines depict the corresponding MVCs. Red points mark the 1000 regions with the largest scaled variances.

### Supplementary Figure2

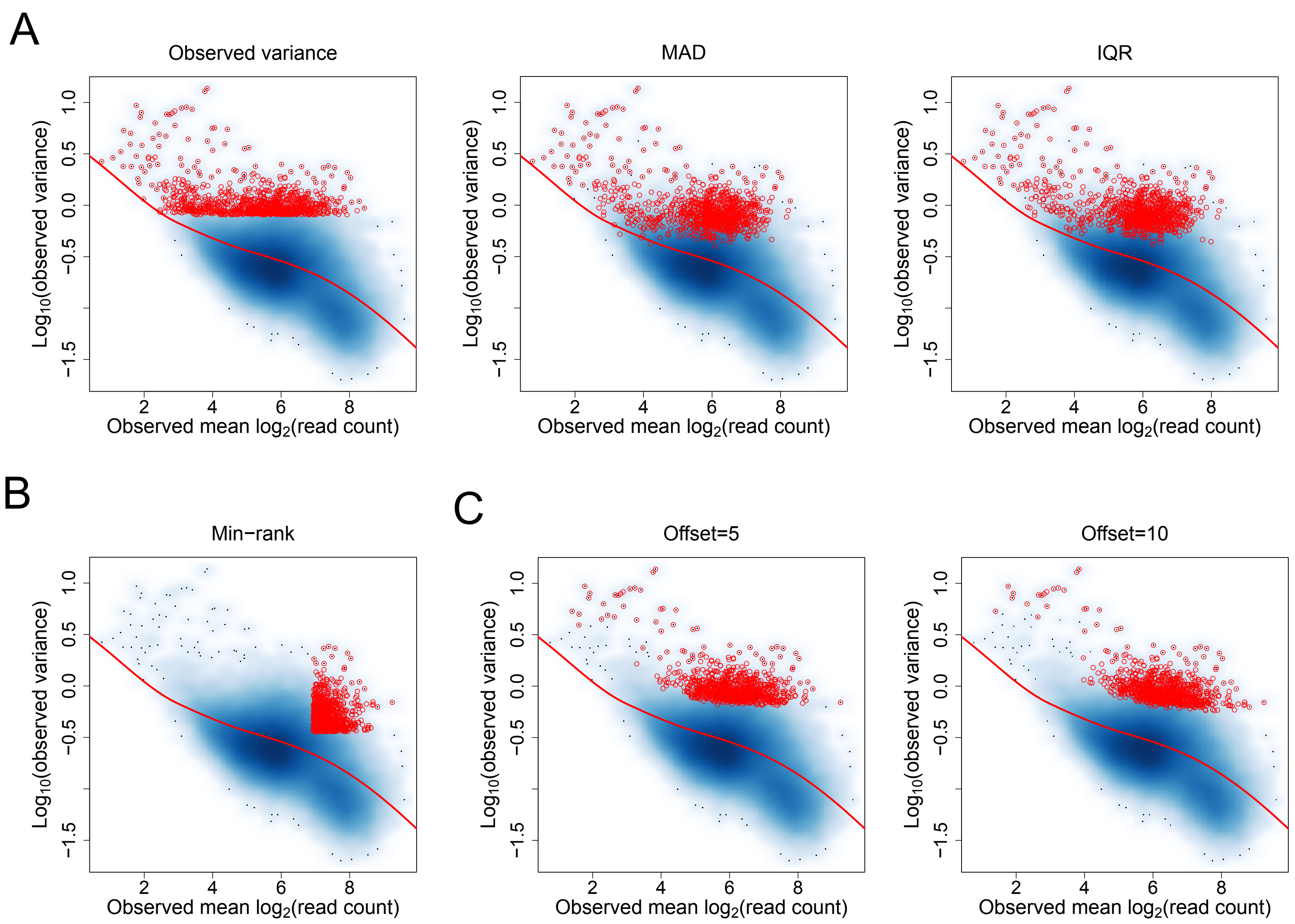

**Supplementary Figure 2. Applying other methods for ranking genomic regions and selecting HVRs. (A-C)** Scatter plots showing the mean-variance trend (at proximal regions) associated with the H3K27ac ChIP-seq data set as well as the regions that are ranked in the top 1000 HVRs by each method (marked by red points). MAD, median absolute deviation; IQR, interquartile range.

### Supplementary Figure3

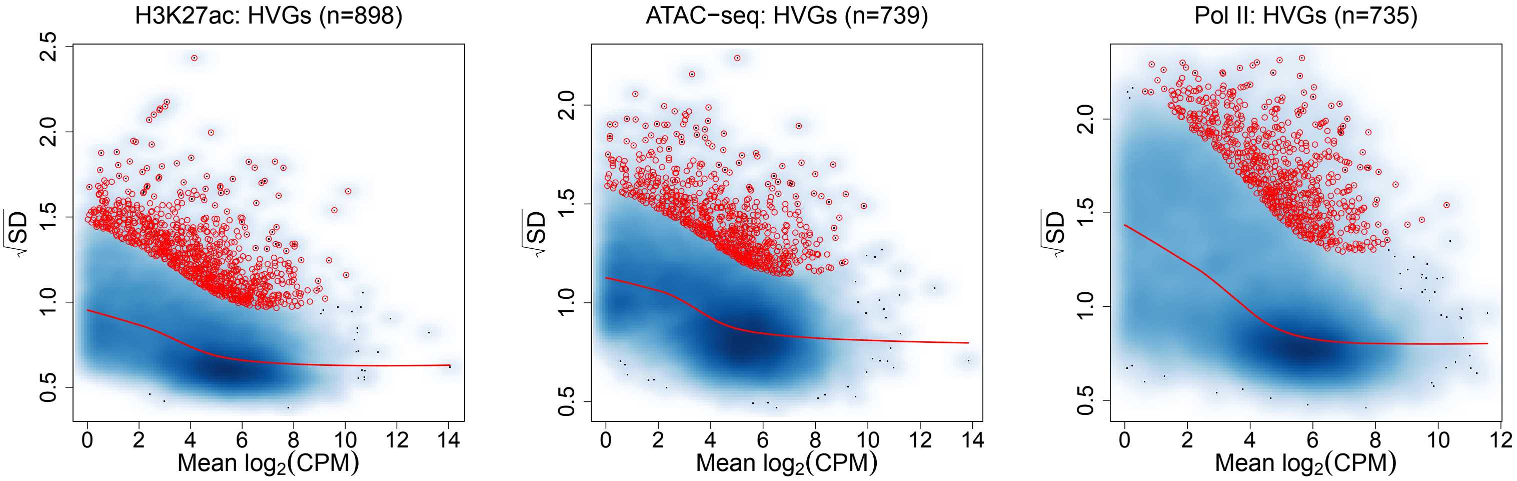

**Supplementary Figure 3. Identifying HVGs.** We have separately identified HVGs for each data set in Table 1, by applying limma-trend to the corresponding RNA-seq data (see Methods in the main text for details). Scatter plots shown here demonstrate the modeling of the mean-variance relationships by limma-trend. Red points in each plot mark the identified HVGs. CPM, count per million; SD, standard deviation.

### Supplementary Figure4

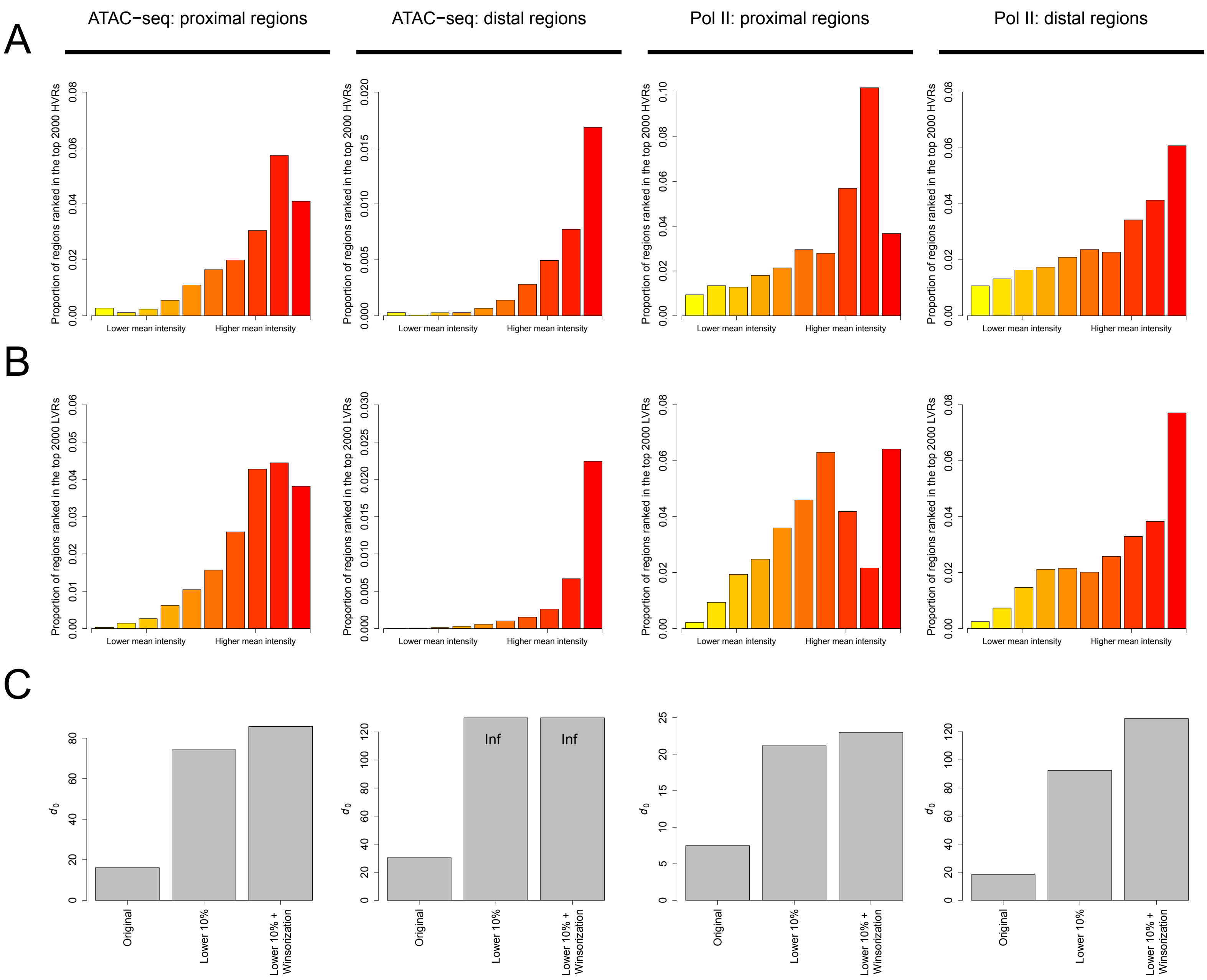

**Supplementary Figure 4. Selecting a subset of genomic regions and using Winsorization for parameter estimation. (A)** For the ATAC-seq and Pol II ChIP-seq data sets, bar plots showing the distributions of top-ranked proximal/distal HVRs along the range of mean intensities. For each data set, proximal and distal regions have been separately divided into 10 equally-sized groups based on the observed mean signal intensities. **(B)** Bar plots showing the distributions of top-ranked proximal/distal LVRs along the range of mean intensities. **(C)** $d_0$ estimates resulting from different parameter estimation methods. Inf refers to positive infinity.

### Supplementary Figure5

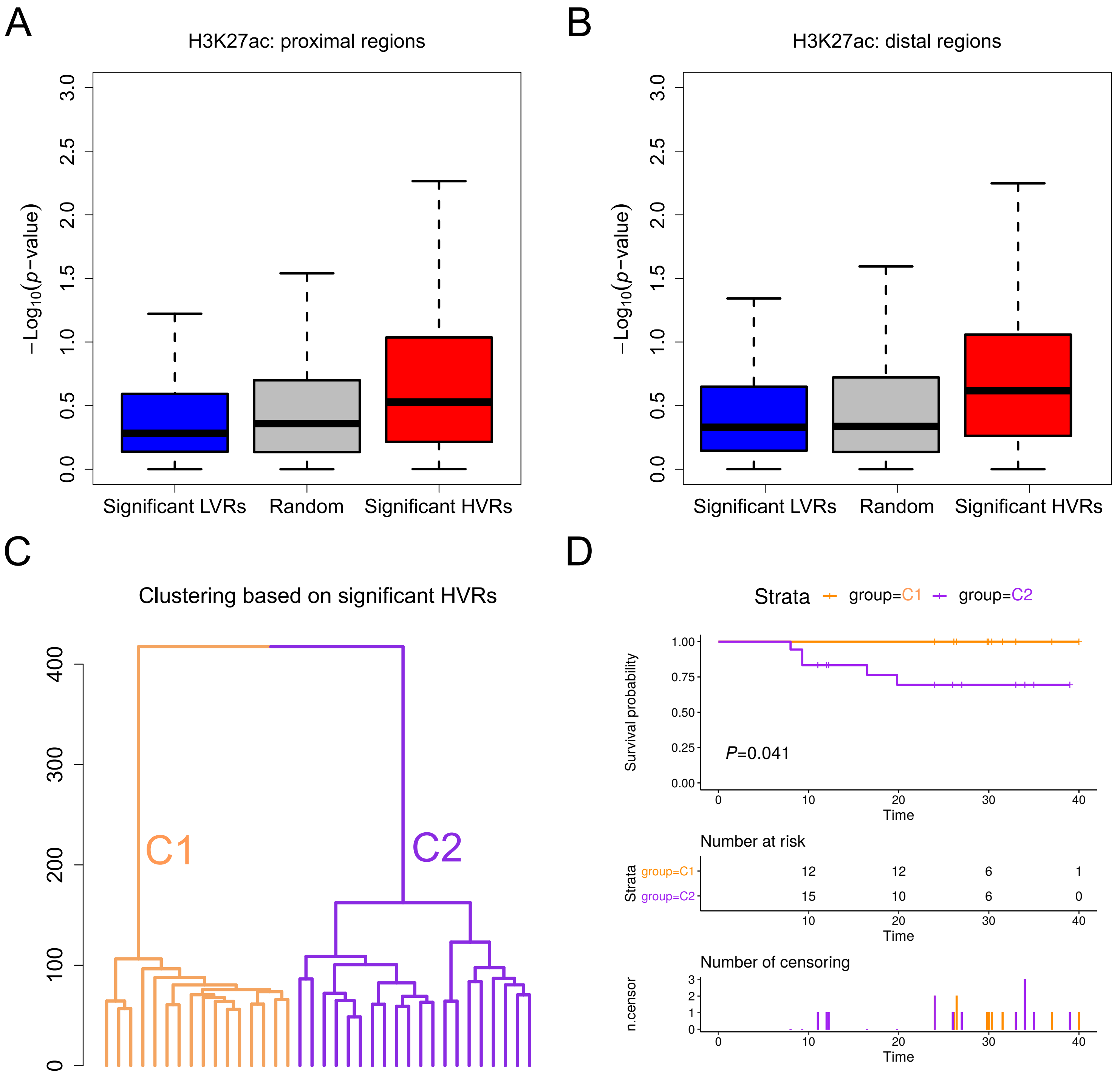

**Supplementary Figure 5. Evaluating the prognostic associations of different genomic regions. (A, B)** Proximal/distal HVRs are more significantly associated with the survival time of patients than proximal/distal LVRs and randomly selected proximal/distal peak regions. Results shown here are based on the H3K27ac ChIP-seq data set. The *p*-values are derived by separately performing a Cox regression on the H3K27ac level in each region. **(C)** Dendrogram showing the hierarchical clustering of the patients based on the proximal and distal HVRs. The patients are classified into two sub-groups, labeled C1 and C2. **(D)** There is a significant survival difference between C1 and C2.

### Supplementary Figure6

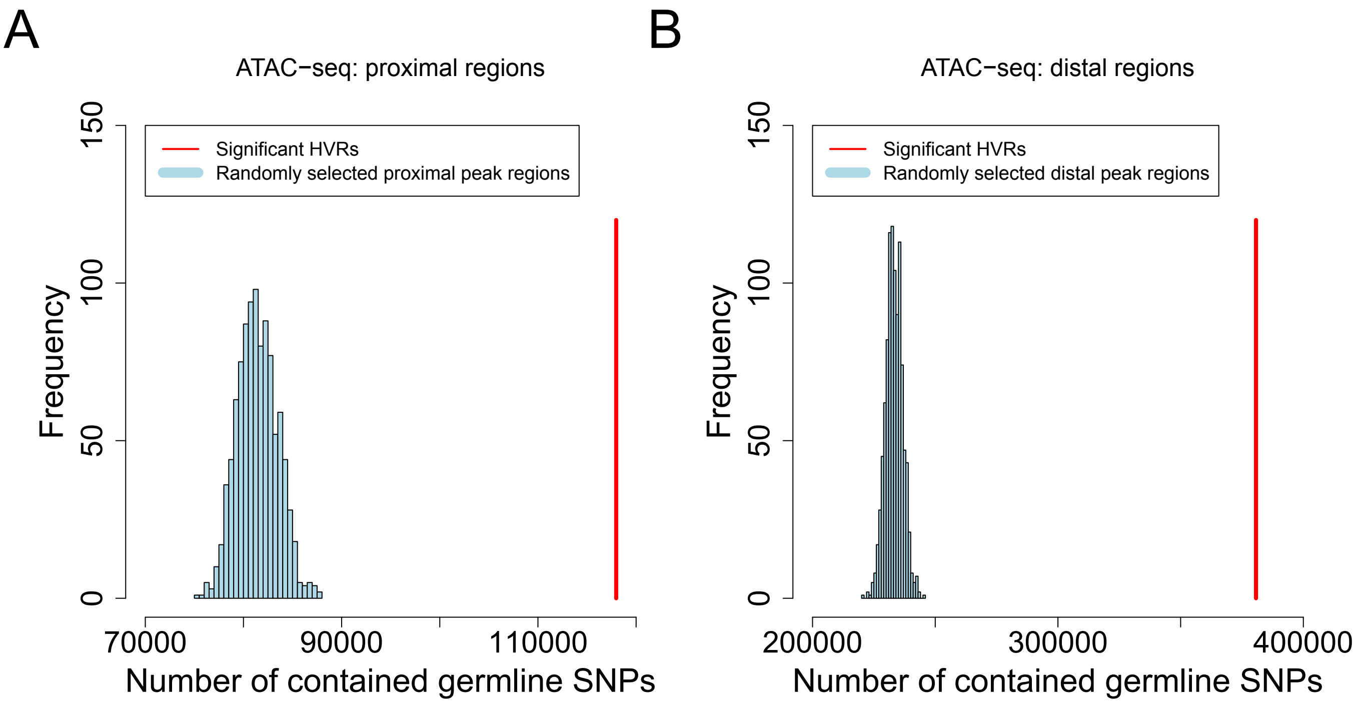

**Supplementary Figure 6. HVRs contain significantly more germline SNPs than by chance. (A)** Significant proximal HVRs defined for the ATAC-seq data set are enriched with germline SNPs. We have performed 1,000 times of random simulation. In each time, a set of proximal peak regions matching the number of the HVRs has been randomly selected. **(B)** Significant distal HVRs are enriched with germline SNPs as well.

### Supplementary Figure7

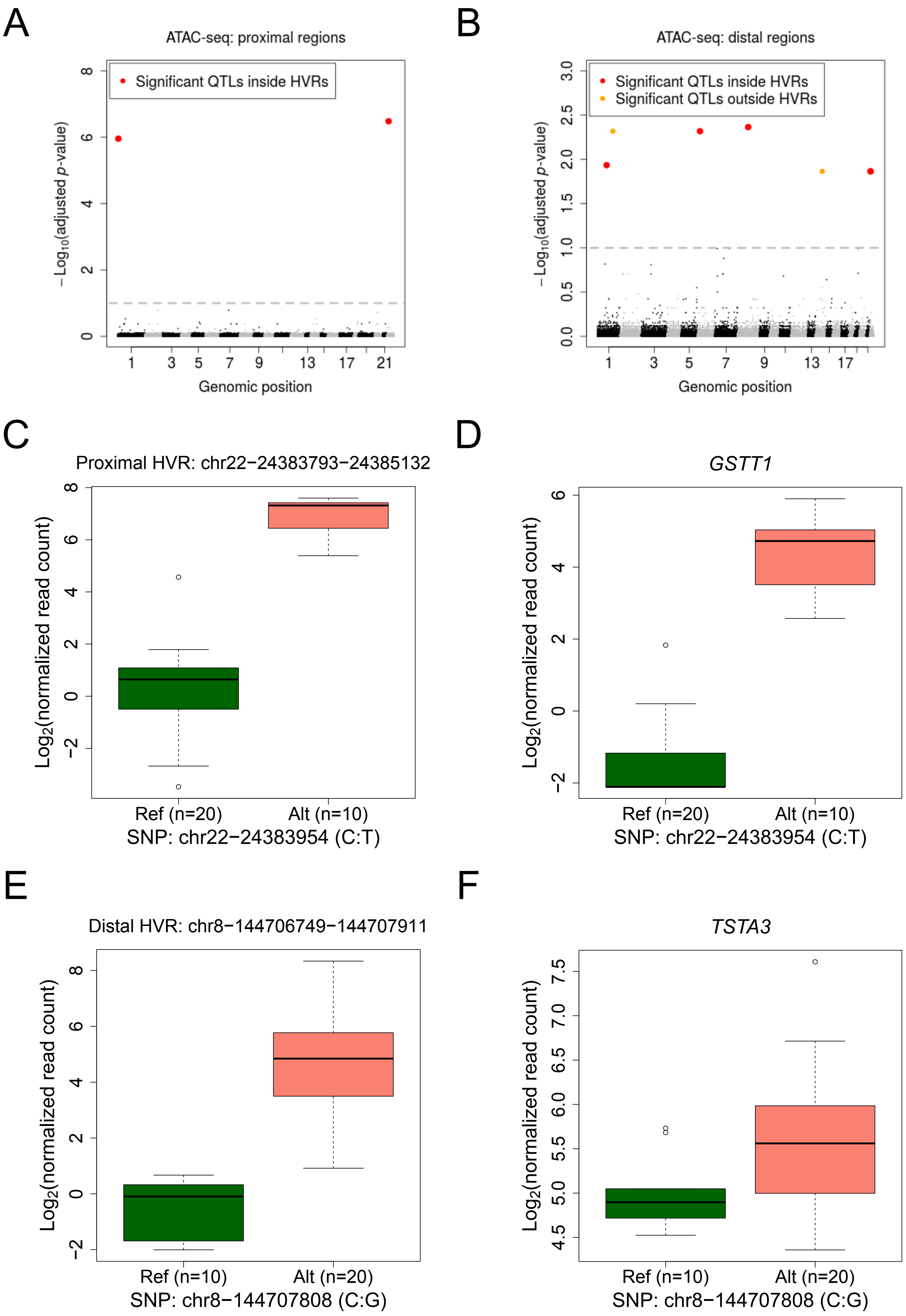

**Supplementary Figure 7. Association between QTLs and HVRs. (A, B)** Identifying QTLs among the germline SNPs located within ATAC-seq peak regions. Each (BH-adjusted) *p*-value assesses the statistical significance of the association between the genotype of a SNP and the ATAC-seq signal in the peak region containing it. **(C)** Box plots showing the ATAC-seq signals associated with different genotypes of the most significant QTL, which is located within a proximal HVR. Ref and Alt refer to the reference genotype and the alternative one, respectively. **(D)** Box plots showing the RNA-seq signals of the downstream gene of the proximal HVR. **(E)** Box plots showing the ATAC-seq signals associated with different genotypes of the most significant distal QTL, which is also located within an HVR. **(F)** Box plots showing the RNA-seq signals of the gene nearest to the distal HVR.

### Supplementary Figure8

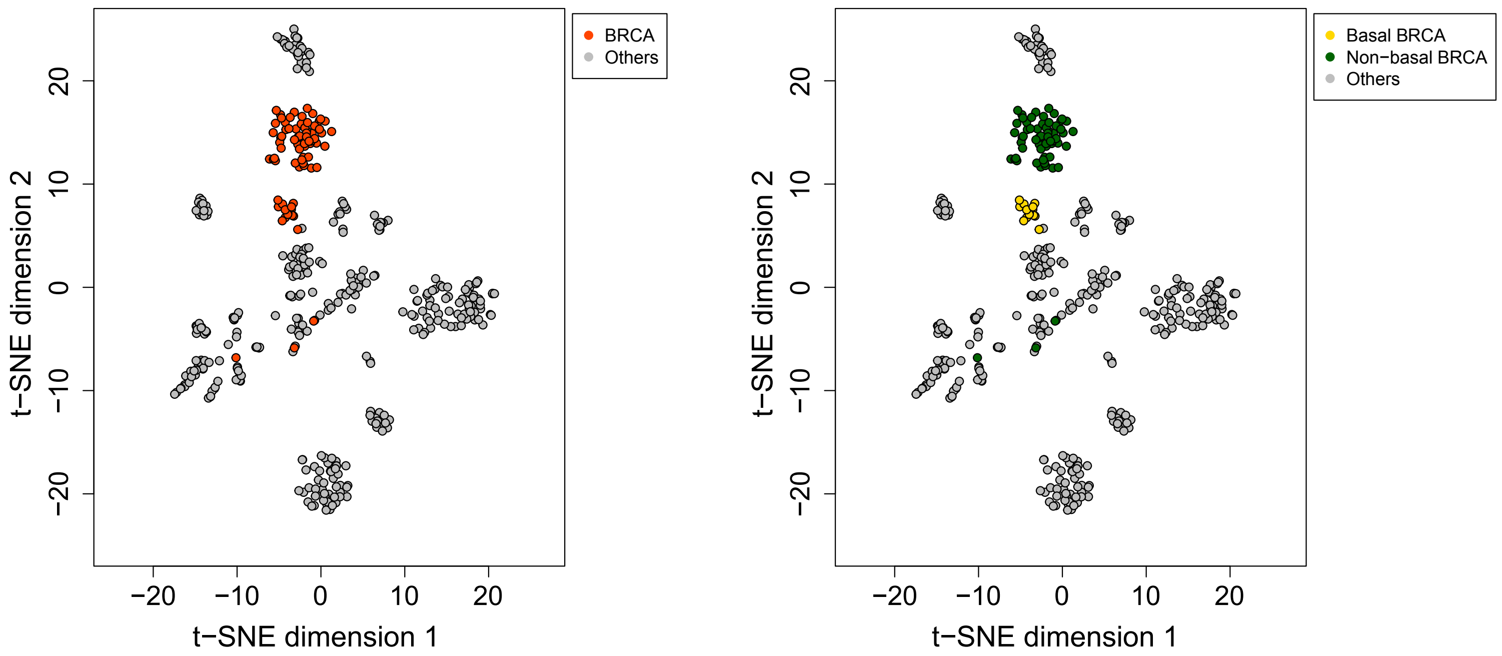

**Supplementary Figure 8. For the TCGA ATAC-seq data set, two-dimensional t-SNE plot showing the distribution of all BRCA patients.** These BRCA patients are comprised of 14 basal and 61 non-basal cases.

### Supplementary Figure9

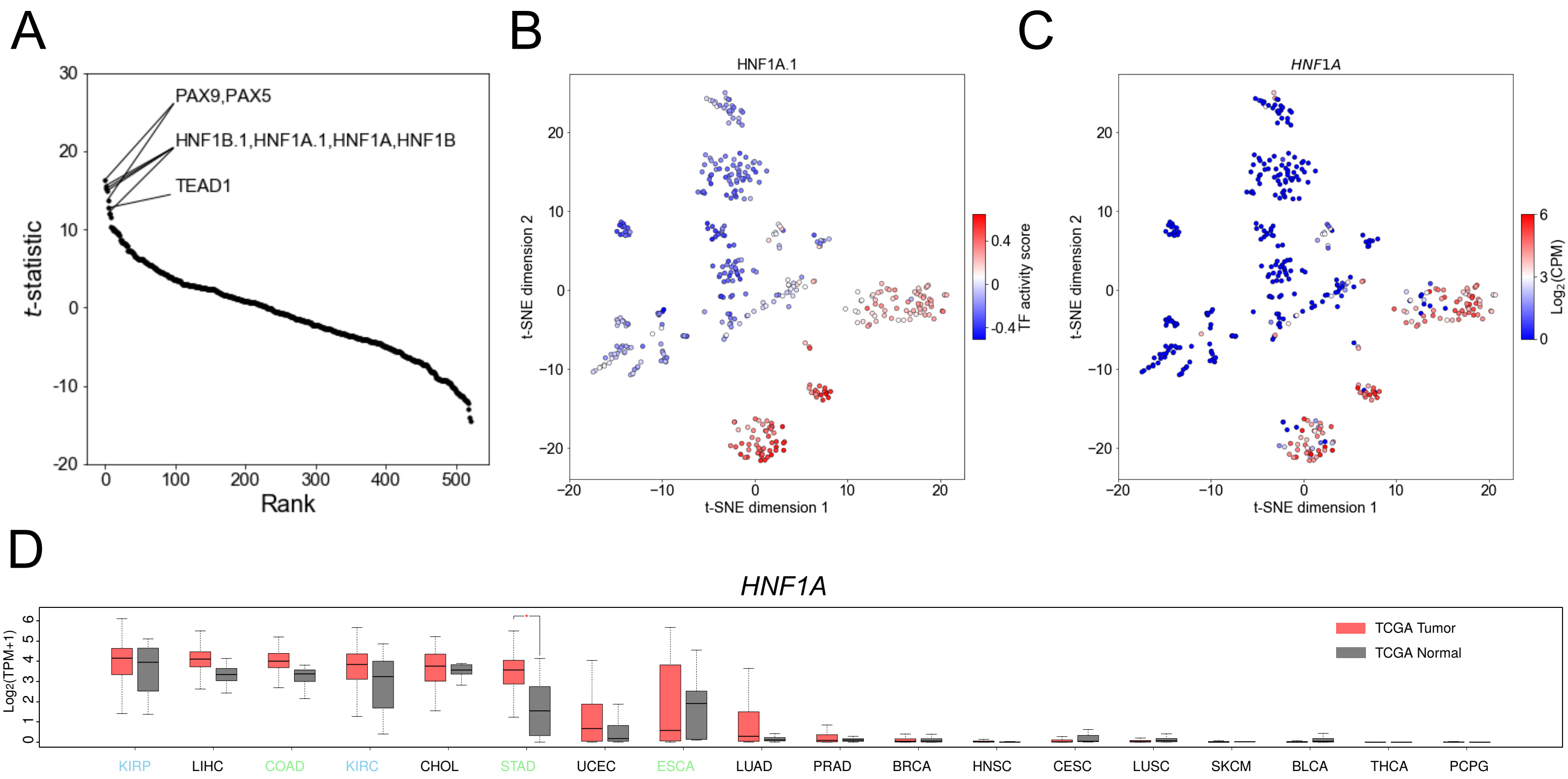

**Supplementary Figure 9. *HNF1A* is identified as a top-ranked TF for the kidney carcinoma class. (A)** Plotting the *t*-statistics of all motifs against their rankings in the identification of TFs specific to the kidney carcinoma class. **(B)** Mapping the TF activity scores associated with the HNF1A.1 motif to the t-SNE plot. **(C)** Mapping the expression levels of the *HNF1A* gene to the t-SNE plot. **(D)** Box plots showing the expression of *HNF1A* in a larger TCGA cohort of patients. TPM, transcripts per million.

### Supplementary Figure10

A

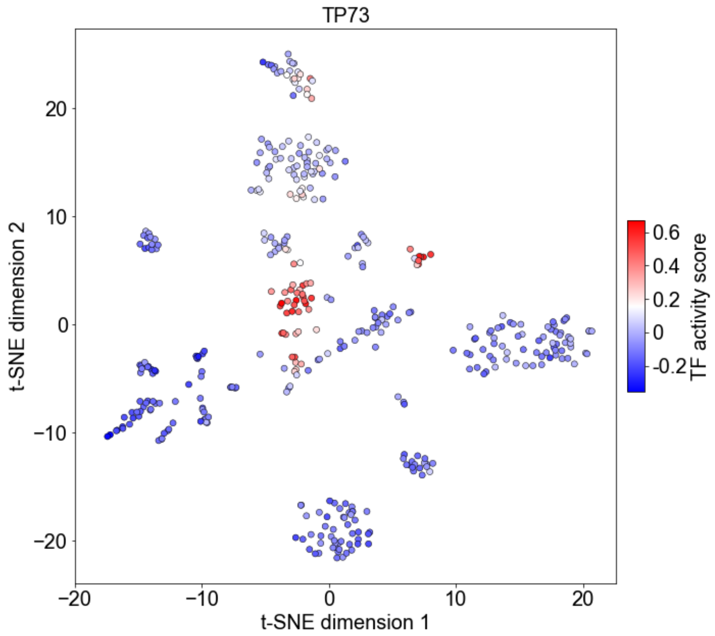


B

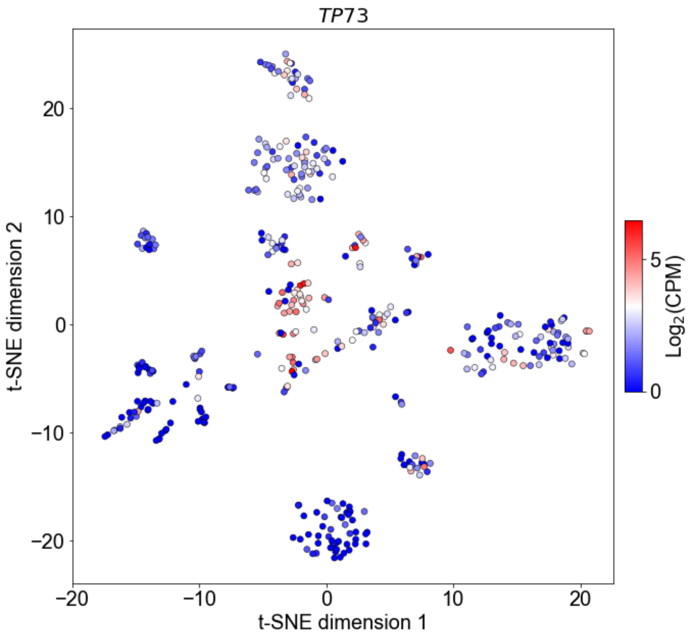


C

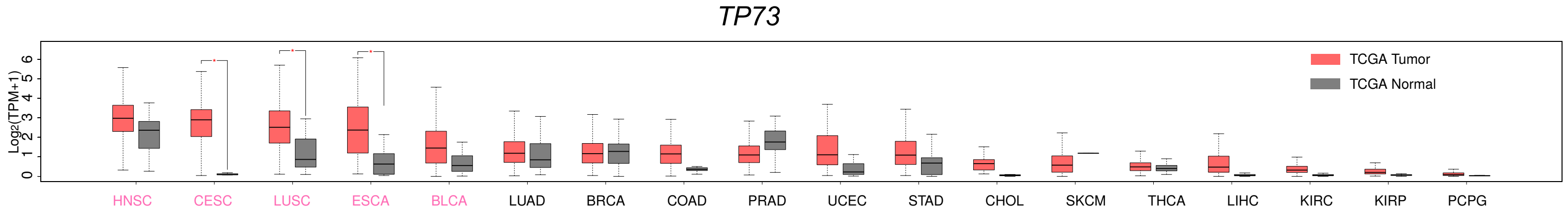

**Supplementary Figure 10.** ***TP73*** **is identified as a top-ranked TF for the SC class.** **(A)** Mapping the TF activity scores associated with the TP73 motif to the t-SNE plot. **(B)** Mapping the expression levels of the *TP73* gene to the t-SNE plot. **(C)** Box plots showing the expression of *TP73* in the larger TCGA cohort.

### Supplementary Figure11

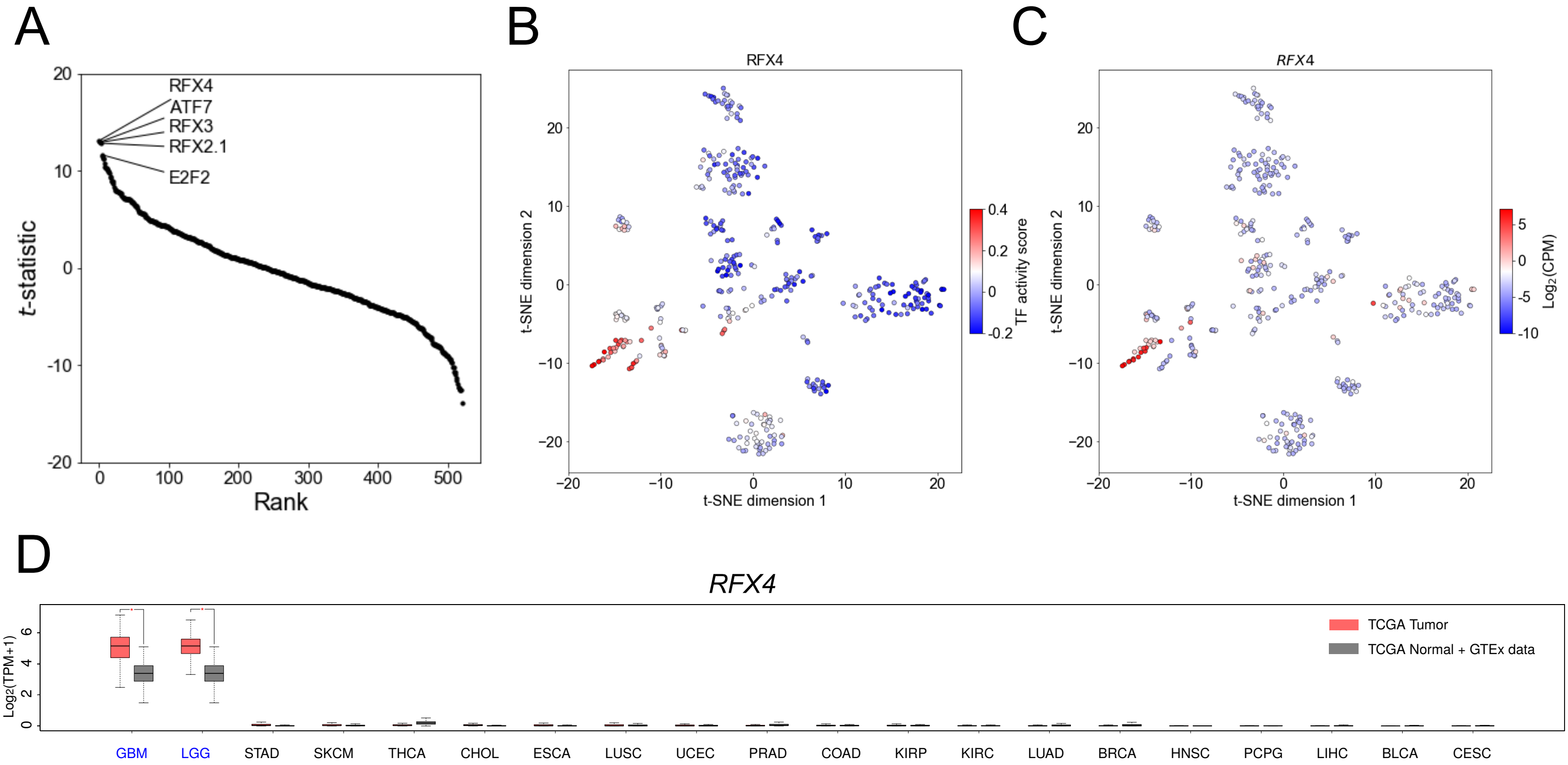

**Supplementary Figure 11. *RFX4* ranks first among the brain cancer class-specific TFs. (A)** Plotting the *t*-statistics of all motifs against their rankings in the identification of TFs specific to the brain cancer class. **(B)** Mapping the TF activity scores associated with the RFX4 motif to the t-SNE plot. **(C)** Mapping the expression levels of the *RFX4* gene to the t-SNE plot. **(D)** Box plots showing the expression of *RFX4* in the larger TCGA cohort as well as in 3,006 RNA-seq samples of normal individuals provided by the GTEx (Genotype-Tissue Expression) project (https://gtexportal.org/home/). We involved the GTEx data because RNA-seq samples for matched normal tissues of GBM and LGG were missing in the TCGA program.
