## Supplementary Table 1-3 for "HyperChIP for identifying hypervariable signals across ChIP/ATAC-seq samples"

| GO term | $-\text{Log}_{10}$(*p*-value) |
|---|---|
| histone lysine demethylation (GO:0070076) | 4.52 |
| tRNA modification (GO:0006400) | 4.13 |
| RNA catabolic process (GO:0006401) | 4.01 |
| histone H3 acetylation (GO:0043966) | 3.90 |
| histone acetylation (GO:0016573) | 3.84 |
| cellular macromolecule catabolic process (GO:0044265) | 3.70 |
| histone H3-K4 demethylation (GO:0034720) | 3.32 |
| energy coupled proton transport, down electrochemical gradient (GO:0015985) | 3.28 |
| cellular response to gamma radiation (GO:0071480) | 3.00 |
| purine ribonucleoside monophosphate biosynthetic process (GO:0009168) | 2.97 |
| DNA metabolic process (GO:0006259) | 2.93 |
| nucleobase-containing compound catabolic process (GO:0034655) | 2.80 |
| purine-containing compound biosynthetic process (GO:0072522) | 2.78 |
| RNA metabolic process (GO:0016070) | 2.72 |
| regulation of generation of precursor metabolites and energy (GO:0043467) | 2.72 |
| genetic imprinting (GO:0071514) | 2.64 |
| NADH metabolic process (GO:0006734) | 2.64 |
| cellular response to ionizing radiation (GO:0071479) | 2.61 |
| cristae formation (GO:0042407) | 2.53 |
| ribonucleoside bisphosphate biosynthetic process (GO:0034030) | 2.49 |

**Supplementary Table 1. Top 20 GO terms enriched from the genes linked with the significant proximal LVRs identified for the H3K27ac ChIP-seq data set.**

| GO term | $-\text{Log}_{10}$($p$-value) |
|---|---|
| positive regulation of transcription from RNA polymerase II promoter (GO:0045944) | 6.75 |
| regulation of transcription from RNA polymerase II promoter (GO:0006357) | 6.62 |
| cartilage development (GO:0051216) | 6.12 |
| epidermis development (GO:0008544) | 6.12 |
| surfactant homeostasis (GO:0043129) | 5.65 |
| chemical homeostasis within a tissue (GO:0048875) | 5.65 |
| positive regulation of transcription, DNA-templated (GO:0045893) | 5.52 |
| kidney development (GO:0001822) | 4.90 |
| negative regulation of multicellular organismal process (GO:0051241) | 4.71 |
| chondrocyte differentiation (GO:0002062) | 4.44 |
| regulation of cartilage development (GO:0061035) | 4.38 |
| renal system development (GO:0072001) | 4.35 |
| lung development (GO:0030324) | 4.18 |
| collecting duct development (GO:0072044) | 3.99 |
| renal absorption (GO:0070293) | 3.86 |
| endochondral bone morphogenesis (GO:0060350) | 3.86 |
| endochondral ossification (GO:0001958) | 3.64 |
| positive regulation of epithelial cell differentiation (GO:0030858) | 3.52 |
| negative regulation of transcription, DNA-templated (GO:0045892) | 3.48 |
| negative regulation of cell differentiation (GO:0045596) | 3.43 |

**Supplementary Table 2. Top 20 GO terms enriched from the genes linked with the significant proximal HVRs identified for the H3K27ac ChIP-seq data set.**

| Cancer type | Abbreviation | Annotation | Cohort size |
|---|---|---|---|
| Adrenocortical carcinoma | ACC | | 9 |
| Bladder Urothelial Carcinoma | BLCA | SC | 10 |
| Breast invasive carcinoma | BRCA | | 75 |
| Cervical squamous cell carcinoma and endocervical adenocarcinoma | CESC | SC | 4 |
| Cholangiocarcinoma | CHOL | | 5 |
| Colon adenocarcinoma | COAD | DIAD | 41 |
| Esophageal carcinoma | ESCA | SC or DIAD | 18 |
| Glioblastoma multiforme | GBM | Brain cancer | 9 |
| Head and Neck squamous cell carcinoma | HNSC | SC | 9 |
| Kidney renal clear cell carcinoma | KIRC | Kidney carcinoma | 16 |
| Kidney renal papillary cell carcinoma | KIRP | Kidney carcinoma | 34 |
| Brain Lower Grade Glioma | LGG | Brain cancer | 13 |
| Liver hepatocellular carcinoma | LIHC | | 17 |
| Lung adenocarcinoma | LUAD | | 22 |
| Lung squamous cell carcinoma | LUSC | SC | 16 |
| Mesothelioma | MESO | | 7 |
| Pheochromocytoma and Paraganglioma | PCPG | | 9 |
| Prostate adenocarcinoma | PRAD | | 26 |
| Skin Cutaneous Melanoma | SKCM | | 13 |
| Stomach adenocarcinoma | STAD | DIAD | 21 |
| Testicular Germ Cell Tumors | TGCT | | 9 |
| Thyroid carcinoma | THCA | | 14 |
| Uterine Corpus Endometrial Carcinoma | UCEC | | 13 |

**Supplementary Table 3. Cancer types involved in the TCGA ATAC-seq data set.** Note that in TCGA studies the abbreviation CESC refers to both cervical squamous cell carcinoma and endocervical adenocarcinoma, but all the 4 CESC patients in this data set belong in the former. Note also that the 18 ESCA patients in this data set consist of 12 ESSC (esophageal squamous cell carcinoma) and 6 ESAD (esophageal adenocarcinoma) cases, which belong in the SC and DIAD classes, respectively. SC, squamous cell carcinoma; DIAD, digestive adenocarcinoma.
